# 150 million years of sustained increase in pterosaur flight efficiency

**DOI:** 10.1101/153957

**Authors:** Chris Venditti, Joanna Baker, Michael J. Benton, Andrew Meade, Stuart Humphries

## Abstract

The long-term accumulation of biodiversity has been punctuated by remarkable evolutionary transitions that allowed organisms to exploit new ecological opportunities, and often resulted in large radiations of species. The Mesozoic flying reptiles – pterosaurs – which dominated the skies for over 150 million years (Myr), were the product of such a transition. The ancestors of pterosaurs were small and likely bipedal early archosaurs, which were certainly well adapted to terrestrial locomotion. More than 220 Myr ago, at some point in the Triassic, pterosaurs took flight and subsequently appear to have become capable and efficient flyers. However, the evolutionary processes that led to this efficiency remain enigmatic. Given the lack of proto-pterosaurs it is difficult to study how flight first evolved in this group, but we can test hypotheses about evolutionary changes to the energetics of locomotion following the transition to flight. Early pterosaurs were challenged by the costs of transitioning between forms of locomotion [1, 2] - from terrestrial to aerial. This imposed a steep energetic hill to climb which means that flight must have provided some offsetting fitness benefits. If the initial transition resulted in a form that was very well adapted to flight we would expect to see no directional change in flight efficiency throughout the history of pterosaurs. Alternatively, the transition may have produced a form that was able to fly but, was not under strong selection for efficiency owing to many benefits conferred by the lack of competition in the novel environment. In the latter case the evolutionary signal of natural selection acting to increase efficiency over millions of years should be detectable. Novel phylogenetic statistical methods and biophysical models combined with information from the fossil record mean we now have the opportunity to test this hypothesis.

To account for phylogenetic uncertainty in our analyses we constructed a Bayesian dated posterior sample of phylogenetic trees for 108 pterosaurs using published character state data [3] (Figure 1A and Supplemental Information). We obtained estimates of mass, projected frontal area, wingspan and wing area for 12 species of pterosaur (Table S1) from Henderson (2010) [4]. We excluded *Quetzalcoatlus northropi* owing to controversy about its size and ability to fly [5]. Using a biophysical model of powered flight (a modified version of Pennycuick’s *Flight* model [6, 7]) we calculate an efficiency of flight index (kg m J^-1^), that is the inverse of the cost of transport, CoT^-1^ (Supplemental Information, see Table S2 for the flight model parameterization). The CoT (J kg^-1^ m^-1^) is the metabolic energy required to move a unit mass a unit distance at the least energetically expensive travel speed.

**Figure 1:**
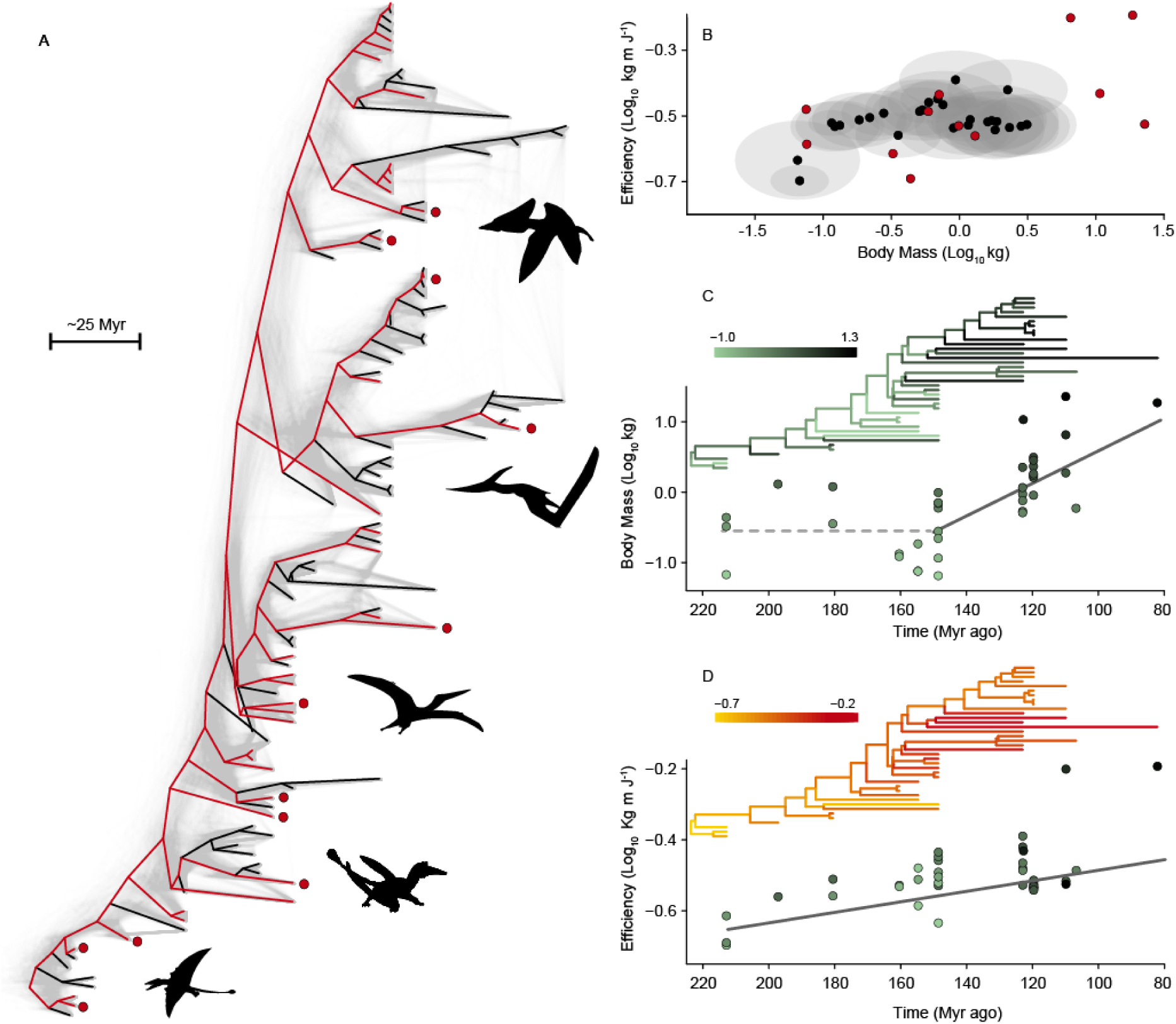
A) A density diagram showing the uncertainty in our phylogenetic reconstruction for pterosaurs (see Supplemental Information for details). Superimposed is the maximum clade credibility tree, where red branches represent phylogenetic coverage of the 38 species we include in our mass and efficiency analyses. At the tips, species with data taken from Henderson (2010) [4] are indicated by red points. B) The relationship between flight efficiency and mass. Our mean phylogenetic imputations are plotted in black and standard deviations are shown by the grey ellipses. As in (A), red points represent species with data [4]. C) The mean predicted phylogenetic regression slopes for our model of mass through time. Points show body mass either as the mean of our posterior estimates or data as taken from Henderson (2010) [4] and are coloured to represent body mass (see inset scale). The phylogenetic tree of our 38 species is coloured using the same scale as the points, such that the colour of each branch represents body mass (imputed for internal nodes from our regression relationship). D) The mean model predictions from our phylogenetic regression for efficiency through time accounting for mass. As in (C), points are either our mean posterior estimates or are taken from Henderson (2010) [4] and are coloured by mass (though are not mass-corrected). Inset is the phylogenetic tree of 38 species where the colour of each branch represents efficiency. For internal nodes, this was imputed using our efficiency through time model.

We used a Bayesian phylogenetic imputation method [8] to produce a posterior distribution of imputed masses, projected frontal area and wing area for a further 26 species of pterosaurs based on phylogenetic regression analyses (Table S1). Phylogenetic regression models were constructed for 12 species with data from Henderson [4] and we used adult wingspan from Benson et al. (2014) [9]. From our imputations we calculated a posterior distribution of 1000 CoT^-1^ estimates for use in our analyses of efficiency through time (Supplemental Information). Our final analyses used information from 38 species that span the majority of the phylogenetic diversity of all known pterosaurs (red branches in Figure 1A).

While CoT^-1^ is an efficiency index related to the amount of energy needed to travel a given distance, independently of how long it takes, we do expect it to be correlated with mass – it is energetically cheaper for a large animal to move a given mass over a particular distance than for a small animal to travel the same distance [10] (Figure 1B). As such, we need to consider mass and is independent evolutionary trajectory in our analyses and interpretation of pterosaur flight efficiency through time. With this in mind, pterosaurs have been reported to conform with Cope’s rule [9] – a well-known phenomenon where species increase in size through geological time. The most compelling evidence for this is derived from analyses reporting an increase in wingspan from ~150 Myr ago to the end of the Cretaceous (~66 Myr ago) coincident with the origin of birds (Avialae) [9]. However, such a trend could emerge as a consequence of increased flight efficiency rather than increase in body size *per se*. Animals with a larger wingspan for their mass are likely to be more efficient flyers [7]. We find, using a phylogenetic regression model that accounts for the uncertainty in our inferred tree and our estimates of species masses (Supplemental Information), that pterosaur size did increase significantly through time. In addition, a model which allows the rate of mass increase through time to differ before and after the origin of the birds fits significantly better. In line with conclusions drawn by Benson et al. [9], we find that there is no significant increase in size until ~150 mya (*Px* = 0.092). From that point the average pterosaur grew significantly (*Px* = 0.005) from 0.25 kg to 9.94 kg, a ~40-fold increase in size, in 65 million years.

Applying our phylogenetic regression to flight efficiency through time we find that, even after accounting for mass, efficiency increased significantly (*Px* = 0.041, Fig 1C). Moreover, in contrast to our finding for mass, there is no significant effect associated with the arrival of birds. Early pterosaurs had an efficiency of 0.23 kg m J^-1^ but by 80 Myr ago they were twice as efficient (CoT^-1^ = 0.43 kg m J^-1^). Our results show that following their transition to aerial locomotion, pterosaurs exhibited a sustained increase in their flight efficiency over 140 million years until their extinction. To achieve this, natural selection acted to decouple the evolution of body size and wing area to sculpt these enigmatic creatures from what might have been inefficient flyers that only took to the air for only short spells, to creatures that could fly long distances over extended periods of time.

Our analyses of how mass changes through time are congruent with other evidence suggesting that the origin of, and competition with, Mesozoic birds had a profound effect on pterosaurs [9], increasingly driving smaller species to extinction. Why the birds had a competitive advantage is not clear, but further investigation into the relative flight efficiency of birds and pterosaurs at small sizes is likely to be enlightening.

Our results highlight that after the appearance of pterosaur flight, there was still significant room for improvement in terms of efficiency. Over millions of years, these animals consistently became better adapted to reduce flight costs - we find no evidence of a slowdown in the rate of efficiency increase (*Px* = 0.747). It is interesting to consider whether this process of fast evolutionary change to a new mode of locomotion, followed by a long but consistent process of fine tuning of efficiency, is a characteristic of all such evolutionary transitions. With that question in mind, our approach demonstrates the power of combining biophysical models and phylogenetic statistical methods with the fossil record to understand the evolution of flight in pterosaurs. In doing so we offer a blueprint to objectively study functional and energetic changes through geological time at a far more nuanced level than has ever before been possible.

